# Redesigned upstream processing enables a 24-hour workflow from *E. coli* cells to cell-free protein synthesis

**DOI:** 10.1101/729699

**Authors:** Max Z. Levine, Byungcheol So, Alissa C. Mullin, Katharine R. Watts, Javin P. Oza

## Abstract

Cell-free protein synthesis (CFPS) platforms have undergone numerous workflow improvements to enable diverse applications in research, biomanufacturing, point-of-care detection, therapeutics, and education using affordable laboratory equipment and reagents. The *Escherichia coli* cell extract-based platform, being one of the most affordable and versatile CFPS platforms, has been broadly adopted. In spite of the promise of simplicity, the cell-free platform remains technically nuanced, posing challenges to reproducible implementation and broad adoption. Additionally, while the CFPS reaction itself can be implemented on-demand, the upstream processing of cells to generate crude cell lysate remains time-intensive, representing one of the largest sources of cost associated with the biotechnology. To circumvent the lengthy and tedious upstream workflow, we have redesigned the processes by developing a long-lasting autoinduction media formulation for cell-free that obviates human intervention between inoculation and harvest. Cell-free autoinduction (CFAI) media supports these advantages through the production of highly robust cell extracts from high cell density cultures nearing stationary phase of growth. Growth of cells to high density and autoinduction of T7 RNAP expression can be achieved by incubation overnight, eliminating the need for user intervention for the entirety of the process. The total mass of cells obtained is substantially increased, which directly results in a 400% increase in total extract volume obtained compared to past workflows. Based on these advances, we outline a new upstream processing workflow that allows researchers to go from cells on a streak plate to completing CFPS reactions within 24 hours while maintaining robust reaction yields of sfGFP (>1 mg/ml). We hope this advance will improve the time and cost-efficiency for existing CFPS researchers, increase the simplicity and reproducibility, and reduce the barrier-to-entry for new researchers interested in implementing CFPS.

## Introduction

Cell-free protein synthesis (CFPS) platforms have provided a robust, flexible, and accessible strategy to express high titers of proteins for the scientific community [1]. The open nature of the platform enables researchers to monitor protein expression in real time, to alter reaction conditions, and to produce traditionally intractable proteins on-demand. CFPS systems have undergone numerous and significant developments over the last 50 years, resulting in long-lived reactions with improved yields at lower costs [2]. The *Escherichia coli-based* CFPS platform in particular has gained traction over the last 30 years and has surpassed the Wheat Germ and Rabbit Reticulocyte platforms in cumulative publications [1]. The broad adoption of the *E. coli*-based crude extracts for CFPS is in part a function of consistent effort by the scientific community to enhance robustness of the platform, streamline the workflow of generating and utilizing cell extracts, and expand the utility and accessibility for new users. From its inception in the 1950s when Nirenberg and Matthaei first used CFPS to decipher the genetic code, there have been numerous advances in both energy systems and laboratory workflows to make CFPS a viable protein expression platform for applications ranging from discovery through manufacturing [3]. Energy systems have been consistently tuned to allow for high protein titers while regenerating substrates to allow for longer lasting reactions with reduced costs [4]. Workflow optimizations include, but are not limited to: growth within baffled flasks, the advancement of sonication-based lysis or bead beating, the utilization of tabletop centrifuges to separate transcriptional and translational machinery from cell lysate, and the ability to scale extract preparation to the 100 L-scale, [5–10]. Most of these advances improved the downstream processing, from cell lysis methods to CFPS reaction conditions to support long-lived, high yielding reactions that are also capable of producing traditionally intractable proteins. The primary improvement to upstream processing over the last 15 years has been the increasing use of baffled shake flasks for cell growth instead of fermenters, otherwise, the process of growing and harvesting cells appears to have remained unchanged [5,11].

Efforts described herein seeks to redefine the upstream processing required to generate *E. coli*-based crude lysates capable of supporting robust CFPS reactions. We define upstream processing as the steps involved in cell growth and harvesting workflow, starting from the originating cell through cell lysis for crude extract preparation. The impetus for improving this workflow is two-fold: A) to reduce the number of technical steps as well as the time and labor associated with upstream processing and B) to improve reproducibility of CFPS from batch-to-batch, user-to-user, and across institutions. The upstream workflow represents the most time-consuming aspect of cell extract preparation, requiring 2-3 days to execute [9,12]. Steps include 1) streak plates from glycerol stocks (day 1); 2) grow seed cultures from streak plates (day 2); 3) inoculate large volume growths with OD_600_ monitoring for induction of T7 RNAP and harvest at precise phases of growth and perform multiple bacterial pellet washing resuspensions prior to storage of cell pellets for later lysis (day 3) [5–7,11]. Downstream processing steps of cell lysis and CFPS reactions can be done immediately following harvest on day three, but often follow on a fourth day that may occur much later in time.

Toward our goal, we have developed a cell-free autoinduction (CFAI) media formulation (Supplemental Table 1) for *E. coli* BL21Star™(DE3) that enables us to obviate the most nuanced and burdensome steps of the existing upstream processing workflow. The outcome is the simplification of a ~3-day workflow down to a 24-hour workflow (Figure 1). Notably, CFAI supports cell growth to high cell densities without sacrificing cell extract productivity. The capacity to generate robust cell extracts from high density cultures results in >400% increase in total extract volume, further improving the value of this approach. Our new approach is simple, reproducible, and decreases the time and labor required, while also increasing the quantity of robust cell extract obtained. Together, the advantages will further reduce the barriers to broad adoption of the CFPS platform.

**Figure 1.**
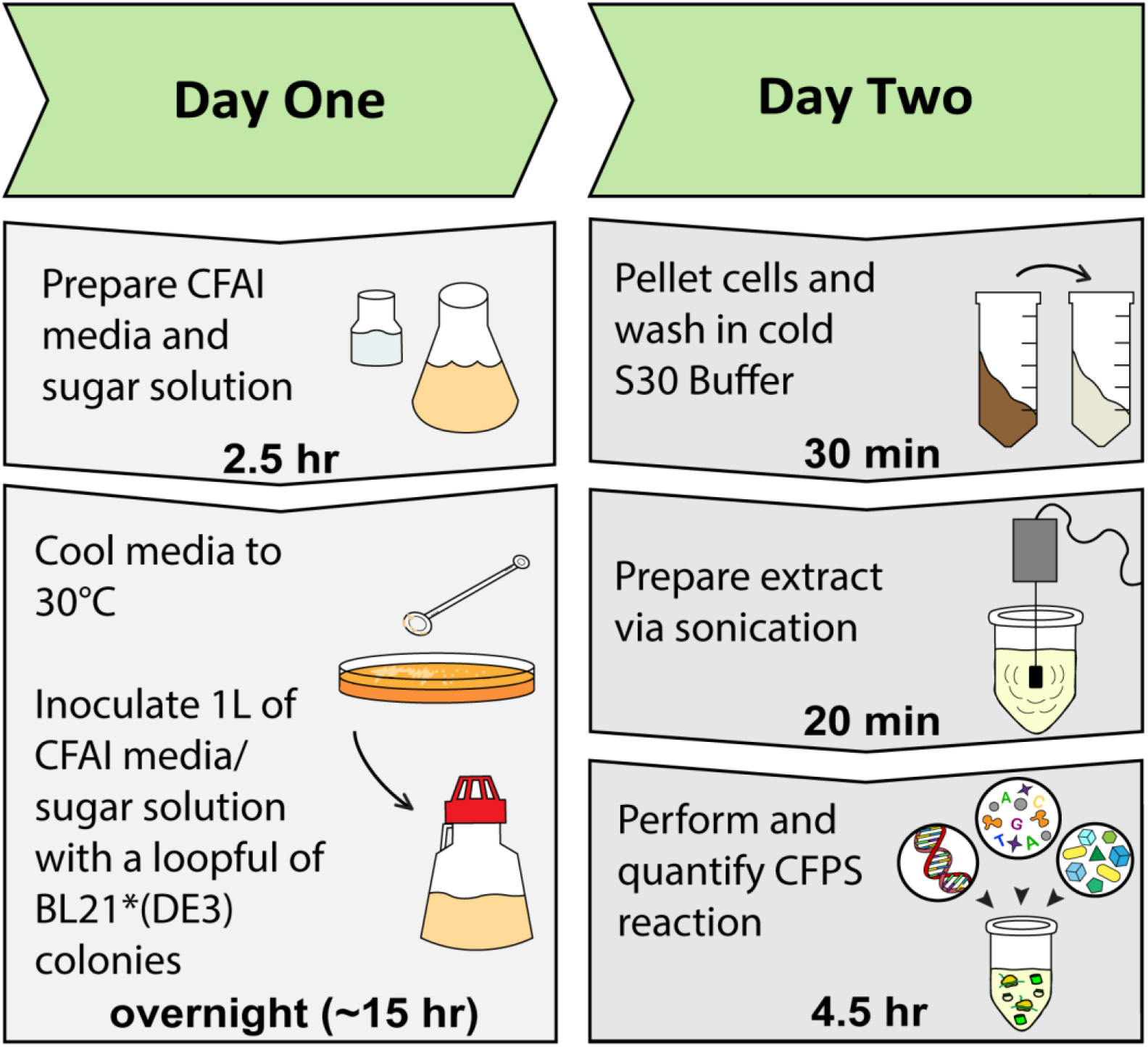
Timeline of CFPS workflow in under 24 hours utilizing the methods presented in this work.

## Methods

### Materials

All materials used in this manuscript have been previously described [9] with the exception of D-lactose (Alfa Aesar), glycerol (Sigma), and MILLEX-HV 0.22 μm Filter Unit (MILLIPORE, Carrigtwohil, Co. Cork, Ireland).

### Cell Growth

All growths derived from *E. coli* BL21Star™(DE3) cells (generously provided by the Jewett Laboratory) are acquired from a glycerol stock and streaked onto a LB agar plate less than two weeks old and stored at 4°C. Streak plates were used within two weeks of inoculation.

#### 2x YTPG Media Growth

A solution of 750 mL 2x YTP was prepared by dissolving 5.0 g sodium chloride, 16.0 g of tryptone, 10.0 g of yeast extract, 7.0 g of potassium phosphate dibasic, and 3.0 g of potassium phosphate monobasic into Nanopure™ water. The solution was adjusted to a pH of 7.2 using 5 M KOH. 250 mL glucose solution was created by combining 250 mL of Nanopure™ water with 18 g of D-glucose. The 2x YTP was transferred to a 2.5 L Tunair™ baffled flask and the glucose solution was transferred to an autoclavable glass bottle. Both solutions were autoclaved for 30 minutes at121°C. A single colony of *E. coli* BL21Star™(DE3) was inoculated into a seed culture of 50 mL of sterile LB and grown overnight at 37°C and 200 rpm. The following day, a 2.5 L Tunair™ baffled flask containing 1 L of 2x YTPG was inoculated from the seed culture to reach an OD_600_ of 0.1. The culture was incubated at 37°C while shaking at 200 rpm until OD_600_ reached 0.6. The 1 L media was then induced with a final concentration of 1 mM of Isopropyl β-D-1-thiogalactopyranoside (IPTG). The culture was then harvested at an OD_600_ of 2.5.

#### AI Media Growth

The autoinduction media was prepared by adopting the recipe developed by Studier, F. W. [13]. In brief, 5.0 g of sodium chloride, 20.0 g of tryptone, 5.0 g of yeast extract, 7.0 g of potassium phosphate dibasic, and 3.0 g of potassium phosphate monobasic were dissolved into 960 mL of Nanopure™ water. The pH was then adjusted to 7.2 using 5.0 M KOH, and autoclaved in the Tunair™ baffled flask for 30 minutes at 121 °C. A separate 40 mL of sugar solution was prepared by dissolving 6.0 mL of 100% glycerol, 2.0 g of D-lactose, and 0.5 g of D-glucose into 34.0 mL of Nanopure™ water. This sugar solution was sterilized using syringe filter sterilization. Following the same procedure for a seed culture, a single colony of E. coli BL21Star™(DE3) was inoculated into 50 mL of LB in a 125-mL Erlenmeyer flask and grown overnight under 37°C at 200 rpm. The next day, a 2.5 L Tunair™ baffled flask containing 1 L of AI media combined with its sugar solution was inoculated by the seed overnight culture to reach an OD_600_ of 0.1. The culture was harvested at an OD_600_ of 2.5.

#### CFAI Media Growth

CFAI media was prepared by dissolving 5.0 g of sodium chloride, 20.0 g of tryptone, 5.0 g of yeast extract, 14.0 g of potassium phosphate dibasic, and 6.0 g of potassium phosphate, monobasic into 960 mL of Nanopure™ water. Subsequently, the pH was adjusted to 7.2 using 5.0 M KOH and autoclaved for 30 minutes at 121 °C. A separate sugar solution was prepared by dissolving 6.0 mL of glycerol, 4.0 g of D-lactose, and 0.5 g of D-glucose into 34.0 mL of Nanopure™ water. The sugar solution was filter-sterilized. The two solutions were mixed to complete the CFAI recipe prior to inoculation. When indicated, glycerol concentrations were titrated (Supplemental Figure 3). The same seed culture inoculation procedure as above was followed for a 2.5 OD_600_ harvest. For high density cultures with no human intervention, a loopful (Supplemental Figure 1) of the previously streaked *E. coli* BL21Star™(DE3) was directly inoculated into 1 L of CFAI media contained in a 2.5 L Tunair™ baffled flask and incubated at 30°C while shaking at 200 rpm. The culture was grown overnight (approximately 15 hours) to an approximate OD_600_ of 10. In some cases, specified amounts of supplemental glycerol were spiked into the culture after overnight growth, an hour prior to harvest.

#### Super CFAI Media Growth

Super-CFAI media consisted of 5.0 g of sodium chloride, 32.0 g of tryptone, 20.0 g of yeast extract, 14.0 g of potassium phosphate dibasic, and 6.0 g of potassium phosphate, monobasic into 960 mL of Nanopure™ water. After the pH was adjusted to 7.2 using 5.0 M KOH, the solution was transferred and autoclaved in a 2.5 L Tunair™ baffled flask and autoclaved for 30 minutes at 121 °C. A separate sugar solution was prepared by dissolving 6.0 mL of glycerol, 4.0 g of D-lactose, and 0.5 g of D-glucose into 34.0 mL of Nanopure™ water. The sugar solution was syringe filter-sterilized. These solutions were combined and inoculated with a loopful of colonies and grown overnight at 30°C shaking at 200 rpm.

#### Cell Harvest

The 1 L media was transferred into a cold 1 L centrifuge bottle (Beckman Coulter, Indianapolis, IN), then centrifuged at 5000 x g and 10°C for 10 minutes (Avanti□ J-E Centrifuge, Beckman Coulter, Indianapolis, IN). After disposing the supernatant, the remaining pellet was transferred to a cold 50 mL Falcon tube using a sterile spatula (SmartSpatula□, LevGo, Inc., Berkeley, CA) while kept on ice. Then, cells were washed once with 40-50 mL of cold S30 buffer (14 mM Mg(OAc)_2_, 10 mM Tris(OAc), 60 mM KOAc, 2 mM dithiothreitol) by resuspension via vortexing with rest periods on ice. In some specified cases, three washes were performed. The resuspension was centrifuged at 5000 x g and 10°C for 10 minutes. After disposing the supernatant, the pellet was weighed, then flash frozen via liquid nitrogen and kept at −80°C until extract preparation. When extracts were prepared during the same day as the harvest, each pellet was flash frozen prior to lysis.

#### Extract Preparation

The frozen cell pellet was combined with 1 mL of S30 buffer per 1 gram of cell pellet and thawed on ice. Once thawed, the cell pellet was resuspended via vortexing with rest periods on ice until no visible clumps of cells were observed. Then, 1.4 mL of the solution was transferred into 1.5mL Eppendorf tubes. A Q125 Sonicator (Qsonica, Newtown, CT) with a 3.175 mm probe was used at a frequency of 20 kHz and 50% amplitude with three forty-five seconds on/fifty-nine seconds off cycles to perform cell lysis. Immediately after, 4.5 μL of 1 M DTT was added to the lysate and inverted several times. The lysate was then centrifuged using a Microfuge 22R Tabletop Centrifuge (Beckman Coulter, Indianapolis, IN) at 18,000 × g and 4 °C for 10 minutes. Following centrifugation, the supernatant was pipetted into a new 1.5 mL Eppendorf tube, flash frozen in liquid nitrogen, and kept in a −80°C freezer until use.

#### Cell-free Protein Synthesis

Cell-free protein synthesis was performed in 15 μL reactions in 1.5 mL Eppendorf tubes in triplicate unless otherwise noted. The standard condition of the reaction included 16 ng/μL of pJL1-sfGFP plasmid, 2.1 μL of Solution A (1.2 mM ATP, 0.850 mM GTP, 0.850 mM UTP, 0.850 mM CTP, 31.50 μg/mL folinic acid, 170.60 μg/mL tRNA, 0.40 mM nicotinamide adenine dinucleotide (NAD), 0.27 mM coenzyme A (CoA), 4.00 mM oxalic acid, 1.00 mM putrescine, 1.50 mM spermidine, and 57.33 mM HEPES buffer), 2.2 μL of Solution B (10 mM Mg(Glu)_2_, 10 mM NH_4_(Glu), 130 mM K(Glu), 2 mM each of the 20 amino acids, and 0.03 M phosphoenolpyruvate (PEP)), 5.0 μL of cell extract, and a varying volume of molecular-grade water to fill the reaction volume to 15 μL [9]. Supplemental reactions included the exogenous addition of 100 μg/mL T7 RNAP (generously provided by the Jewett Laboratory). The cell-free protein synthesis reaction was carried out at 37°C for a minimum of four hours.

#### Quantification of Reporter Protein

Fluorescence intensity of superfolder GFP (sfGFP) was measured in triplicate per reaction with excitation and emission wavelengths of 485 and 528 nm respectively using a half area 96-well black polystyrene plate (Corning Incorporated, Corning, NY) containing 48 μL of 0.05 M HEPES solution (pH 7.2) and 2 μL of the cell-free protein synthesis reaction product. Fluorescence measurements were conducted using a Cytation 5 imaging reader (BioTek, Winwooski, VT). The fluorescence was then converted to concentration of sfGFP (μg/mL) based upon a standard curve as previously described [9].

## Results

In efforts to reduce the time and labor associated with obtaining cells for extract preparations, we first assessed whether three wash cycles of the bacterial pellet were necessary prior to lysis of the cells. We determined that performing one wash instead of three is not detrimental to the resulting cell extracts’ capacity to express the reporter protein sfGFP (Supplemental Figure 2) [5,6]. From this point onward, each cell pellet underwent only one wash regardless of media type. Additionally, we did not perform a runoff reaction as it is not necessary for the BL21Star™(DE3) strain [5].

Next, an autoinduction strategy was employed to obviate the need to induce cells with IPTG, the costly lactose analog. An autoinduction media recipe adopted from F.W. Studier [13] is similar to 2x YTPG in yeast extract, tryptone, and phosphate quantities, but differs significantly in carbon sources, significantly reducing the amount of glucose in favor of added lactose and glycerol (Supplemental Table 1). Replacement of glucose with lactose and glycerol as carbon sources was of concern given that glucose supplementation in 2x YTP was first developed to limit the expression of alkaline and hexose phosphatases that would normally result in a buildup of inorganic phosphates, metabolites detrimental to the CFPS reaction as well as to activate central metabolism for energy recycling within the CFPS reactions [14,15]. To our surprise, replacing 2x YTPG with autoinduction media displayed no significant difference in the extract’s capacity to perform *in vitro* transcription and translation when cells were harvested at an OD_600_ of 2.5 (Figure 2). Reducing the requirement to monitor OD_600_ for T7 RNAP induction and performing 1 wash instead of 3 washes provides minor but noteworthy improvements to the workflow. However, growth in autoinduction media remains dependent on harvesting cells within a precise window of cell growth during the early to mid-logarithmic phase in which cells are undergoing rapid doubling at which point ribosomes and associated translational proteins are thought to be in high abundance [5,16–18]. A downside to this approach is that it tethers the researcher to monitoring cell densities for the duration of the growth, increasing the labor and opportunity cost associated with obtaining cells for CFPS. We sought to test the previous observations that established the optimal OD_600_ for harvesting cells [5,8]. Toward this end, cells were grown to high densities, OD_600_ of 10, in both 2x YTPG and autoinduction media. Our observations confirmed previous findings that extracts generated from cells harvested at high cell densities nearing stationary phase of growth in either 2x YTPG or autoinduction media show a depressed capacity for protein production compared to extracts generated from cells harvested at an OD_600_ of 2.5 (Figure 2).

**Figure 2.**
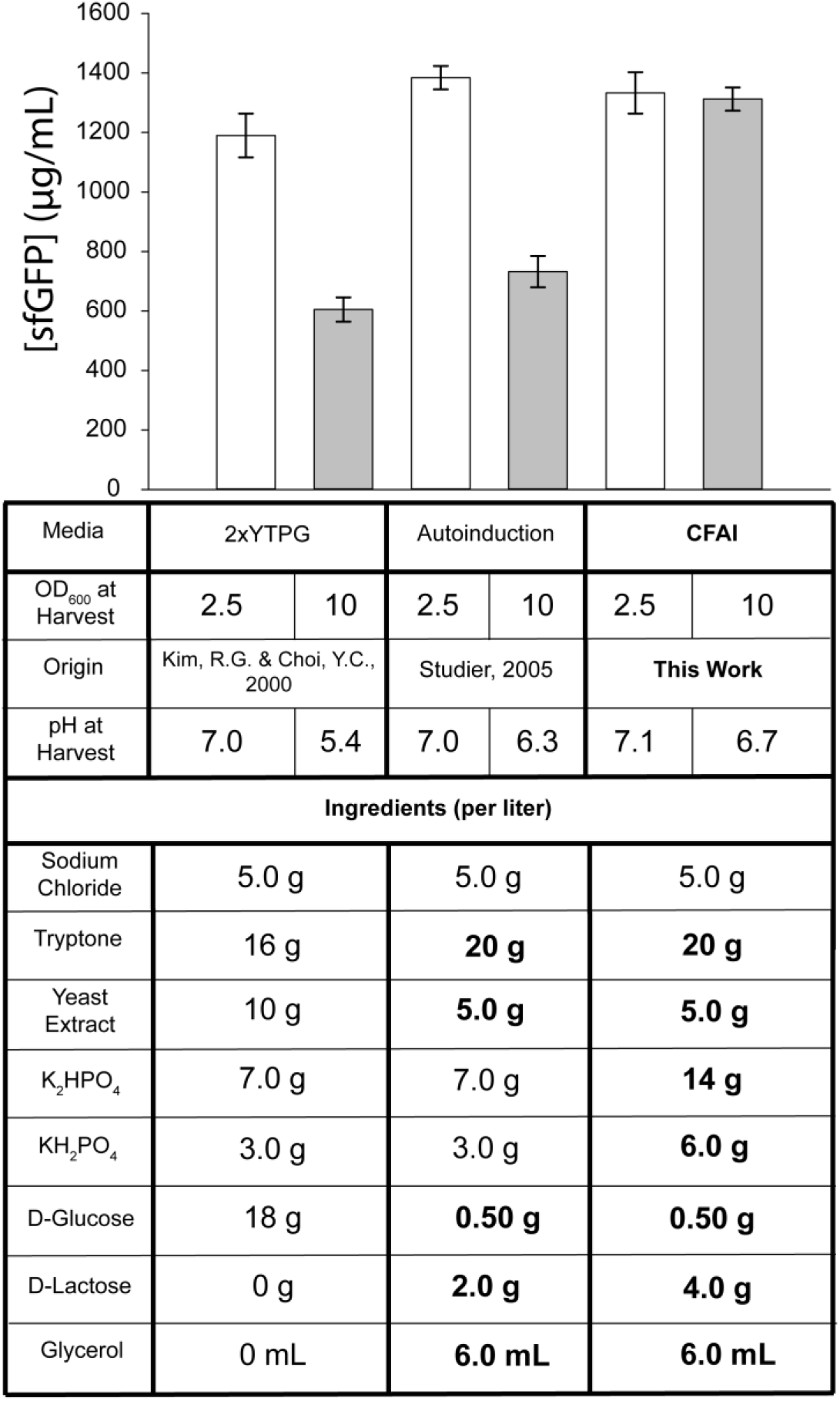
[sfGFP] versus various media recipes harvested at OD_600_ of 2.5 and 10.0. All values are derived from three independent cell extract preparations from three independent 1 L media growths for each condition. Concentration values were calculated from the average of cell-free protein synthesis reactions performed in triplicate for each cell extract that underwent three independent measurements. All error bars represent one standard deviation of the average of three independent reactions for each condition performed in triplicate. Bolded ingredients represent modifications from the 2x YTPG media formulation.

We sought to identify whether the necessity to harvest cells at mid-log phase of growth is a result of functional limitations other than translation machinery. We observed that depressed CFPS yields from high cell density cultures correlate with more acidic culture conditions at harvest (Table 1). To test the role of pH destabilization, we increased the buffering capacity of the AI media by two-fold. Additionally, we hypothesized that the extended growth times may exhaust the lactose carbon source available in the AI media, resulting in depressed expression of T7 RNAP and/or altered metabolism of the cells, becoming incompatible with the PANOxSP energy system in our CFPS reactions. To address these concerns, we also increased the lactose concentration by two-fold within the AI media (Supplemental Table 1). Cells grown in the new media formulation were first cultured to an OD_600_ of 2.5 in order to establish whether the added buffering capacity or lactose are detrimental to the resulting extract. Data displayed in Figure 2 deemed that the extract resulting from the modified AI media performed robustly, yielding >1 mg/mL of reporter protein. Cells were then grown to an OD_600_ of 10 in the high lactose and high buffering capacity autoinduction media, washed once, and processed for extract preparation. The extract resulting from cells grown to high densities resulted in highly active cell extracts capable of producing >1 mg/mL of reporter protein sfGFP (Figure 2). These findings demonstrated that our cell-free autoinduction (CFAI) media formulation expands the limits of the traditional cell growth workflow.

**Table 1.** Cell pellet mass and extract volume generated from corresponding media types grown in triplicate. Values were averaged across triplicate growths.

|  | 2xYTPG |  | AI |  | CFAI |  |
| --- | --- | --- | --- | --- | --- | --- |
| OD<sub>600</sub> | 2.5 | 10 | 2.5 | 10 | 2.5 | 10 |
| pH at Harvest | 7.0 | 5.4 | 7.0 | 6.3 | 7.1 | 6.7 |
| Cell Pellet (g) | 4.21 | 14.22 | 4.79 | 17.4 | 4.13 | 13.9 |
| Extract Volume (mL) | 4.29 | 17.16 | 5.13 | 20.52 | 4.41 | 17.64 |

In order to maximize the potential of CFAI media, we evaluated the optimal concentration of each component of our CFAI media. Toward this end, we tuned the carbon source concentrations and timings of supplementing carbon source, as well as yeast extract and tryptone quantities. Increased concentrations of glycerol were added to the sugar recipe but provided no boost to the overall cell density or extract productivity (Supplemental Figure 3). To test the hypothesis that metabolic shifts as cells approach stationary phase play a role in limiting extract productivity, we also tested conditions where glycerol was spiked into high density cultures 1 hour prior to harvest in efforts to reactivate metabolism. These interventions also did not improve overall cell density or extract productivity compared to CFAI media, confirming that the optimal conditions require minimal human intervention in the workflow. We chose to maintain the current concentration of glucose in order to provide the adequate threshold of energy in the media to begin expressing the enzymes needed to uptake and begin metabolizing lactose [13,19]. To identify whether the full potential of CFAI was limited in other resources, yeast extract and tryptone were also augmented based on the SuperBroth media recipe that is marketed for high density cultivation of *E. coli* cells [20]. The resulting Super-CFAI media displayed similar OD_600_ values and extract productivity levels as the CFAI media. These findings suggest that the added cost of reagents for the Super CFAI media are not justified and that the CFAI media formulation is optimal (Supplemental Figure 4).

The capacity to obtain highly productive cell extracts from high density cell cultures using CFAI media liberates the researcher from the time and labor associated with existing workflows for cell growth. To expand on this capacity, we sought to reduce or remove human intervention from all cell growth steps involved in the upstream processing. Specifically, the traditional workflow requires the researcher to 1) generate streak plates, 2) inoculate seed cultures from the colonies grown on the streak plates, 3) inoculate larger volumes of media with the seed culture cells, and 4) monitor growth of cells that will ultimately generate cell extract capable of *in vitro* transcription and translation. Given the slow nature of cell propagation, this process consumes 2-3 days. We tested a modified workflow in which colonies (Supplemental Figure 1) of BL21Star™(DE3) from a streak plate were inoculated directly into 1 L of CFAI media, incubated for 15 hours overnight, and harvested the subsequent morning. This experiment was conducted at both 30°C and 37°C, and the resulting OD_600_ values were 8.0 and 10.0 respectively, generating cell pellets of 15 g and 18 g respectively. Cells were washed once during harvest and lysed via sonication for extract preparation. Extracts from both overnight growths were robust, yielding >1 mg/mL of sfGFP, with the 30°C growth producing a ~10% higher titer than the 37°C growth. If streak plates and CFAI media are available, this new workflow enables researchers to inoculate a liquid culture at 5 p.m., harvest at 8 a.m., generate extracts by 10 a.m., setup CFPS reactions by noon, and quantify by 3-4 p.m. In other words, this new workflow enables researchers to go from cells on a streak plate to conducting and analyzing CFPS within 24 hours with under 6 hours of a researcher’s active effort.

CFAI-based high density cell growth provides advantages beyond improved workflows. The quantity of cells obtained from high density growths are ~4 times greater, and the corresponding extract volumes obtained are also ~4 times larger (Table 1). As a function of the simplicity, the CFAI-based workflow is also highly reproducible. To evaluate this, we grew three independent cultures of each condition, performed three independent extract preparations of each growth, tested each extract in triplicate CFPS reactions, and subsequently quantified productivity of each reaction in triplicate. The standard deviation resulting from these independent replicates is under 10% (Figure 2) underscoring the reproducibility of the approach. Lastly, while the cost of 2x YTPG and CFAI media are similar, increased extract volumes, combined with reduced researcher time, makes this new approach significantly more cost-effective.

## Discussion

The results presented here demonstrate the development of a new upstream workflow for performing *E. coli* crude lysate-based cell-free protein synthesis. The new approach provides four key advantages over past workflows by: 1) decreasing the overall time by from a four day process to just under 24 hours, 2) decreasing the labor and oversight required from the researcher, 3) increasing the extract obtained by ~400%, 4) removing the need to introduce exogenous T7 RNAP to CFPS reactions. Directly inoculating a 1 L volume of media with a loopful of colonies and obviating the seed culture reduces the workflow by an entire day’s time. Although standard microbiology growth procedures often rely on a single colony to limit genetic diversity, the streak plate is generated from an isogenic glycerol stock. Additionally, many biotechnology endeavors utilize the inoculation of multiple colonies into a liquid cultures to support their biotechnology applications [21,22]. Moreover, the cell extracts produced from our growths have been shown to have reproducible robustness from batch-to-batch, reducing immediate concerns associated with the genetic diversity arising from multiple colonies. For these reasons, we maintain that inoculating with multiple colonies from a fresh plate (less than two weeks old and stored at 4°C) of BL21Star™(DE3) that is generated from an isogenic glycerol stock is suitable for CFPS applications. Next, the rationale for using a seed culture is to expedite cell growth in large volumes; the seed culture allowed researchers to begin growth of a 1 L culture at an OD_600_ of 0.1 - 0.3 in order to reach OD_600_ of 2.5 in a timely manner. The capacity to obtain robust extracts from high density cell cultures that have autoinduced T7 RNAP expression obviates the need to monitor cell densities for induction of T7 RNAP between OD_600_ of 0.6 - 0.8 or for harvest at mid-log phase and therefore, eliminates the need for seed cultures [5–7,9,11].

Following cell harvest, the time needed to wash the bacterial pellet is reduced to a third by using one washing step instead of three which displayed no drop in overall productivity of the cell extract (Supplemental Figure 2). Given that cell pellets are increasingly difficult to resuspend after each wash, the practical time and labor savings are likely greater than 3-fold. Moreover, our recipe still allows for a typical OD_600_ 2.5 harvest if large amounts of extract are not necessary, and it achieves this cell density at a faster growth rate than standard 2x YTPG media (Supplementary Figure 5). As a result, a researcher can inoculate a loopful of colonies in the morning and harvest at OD_600_ 2.5 seven hours later prior to going home for the day. In addition to the aforementioned advantages, researchers looking to maintain their current workflows may find CFAI superior to 2x YTPG for improved growth rates as another source of time reduction. Lastly, our data showed that CFAI-based extracts are not limited by T7 RNAP and do not benefit from supplementation of purified enzyme which suggests that there is sufficient induction of the *lac* operon throughout the growth period (Supplemental Figure 6).

In all, these efforts have resulted in the development of a new upstream workflow for the preparation of *E. coli* extract. The CFAI media-based workflow provides researchers with an economical and reproducible strategy to generate large volumes of robust cell extracts capable of producing over 1 mg/mL of reporter protein. Notably, a researcher stocked with CFAI media and a streak plate can go from cells to CFPS within 24 hours in a ‘set it and forget it’ manner. We hope this innovation will transform the workflow for existing CFPS researchers and reduce the barrier to entry for new users.

## Acknowledgements

Authors would like to acknowledge Dr. Jennifer VanderKelen and Andrea Laubscher for technical support, Wesley Kao, Nicole Gregorio, Phillip Smith, and Logan Burrington for helpful discussions. Authors also acknowledge funding support from the Bill and Linda Frost Fund, Center for Applications in Biotechnology’s Chevron Biotechnology Applied Research Endowment Grant, Cal Poly Research, Scholarly, and Creative Activities Grant Program (RSCA 2017), and the National Science Foundation (NSF-1708919). MZL would like to acknowledge the California State University Graduate Grant.

## Disclosures

The authors declare that they have no competing financial interests or other conflicts of interest.

