## Supplemental Information for "Redesigned upstream processing enables a 24-hour workflow from *E. coli* cells to cell-free protein synthesis"

Autoclave for 30 minutes, 121°C:

| | 2x YTPG Media | AI Media | CFAI Media | Super-CFAI Media |
|---|---|---|---|---|
| Sodium Chloride | 5.0 g | 5.0 g | 5.0 g | 5.0 g |
| Tryptone | 16.0 g | 20. g | 20. g | 32 g |
| Yeast Extract | 10.0 g | 5.0 g | 5.0 g | 20.0 g |
| Potassium phosphate, dibasic | 7.0 g | 7 .0 g | 14. g | 14 g |
| Potassium phosphate, monobasic | 3.0 g | 3.0 g | 6.0 g | 6.0 g |
| Nanopure™ Water | Up to 750 mL | Up to 960 mL | Up to 960 mL | Up to 960 mL |

Filter Sterilize:

| | 2x YTPG Media* | AI Media | CFAI Media | Super-CFAI Media |
|---|---|---|---|---|
| D-Glucose | 18.0 g | 0.50 g | 0.50 g | 0.50 g |
| D-Lactose | 0 g | 2.0 g | 4.0 g | 4.0 g |
| Glycerol | 0 mL | 6.0 mL | 6.0 mL | 6.0 mL |
| Nanopure™ Water | Up to 250 mL | Up to 40 mL | Up to 40 mL | Up to 40 mL |

**Supplemental Table 1**. Ingredient recipes for various media and sugar solutions used to make 1 L of media in this work. (* sugar solution that may undergo autoclaving for 30 minutes, 121°C).

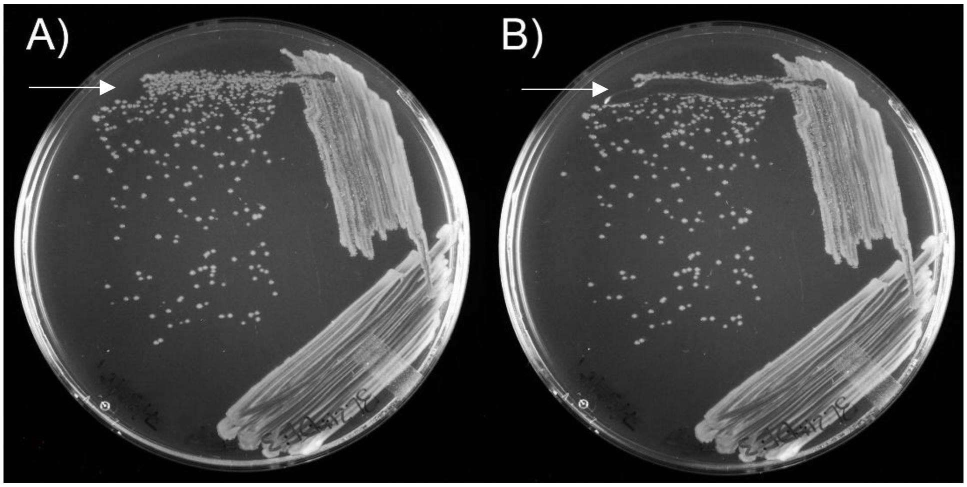


**Supplemental Figure 1**. Pictures of a BL21*(DE3) LB streak plate before (A) and after (B) removing a loopful of colonies for direct inoculation into 1 L of media.

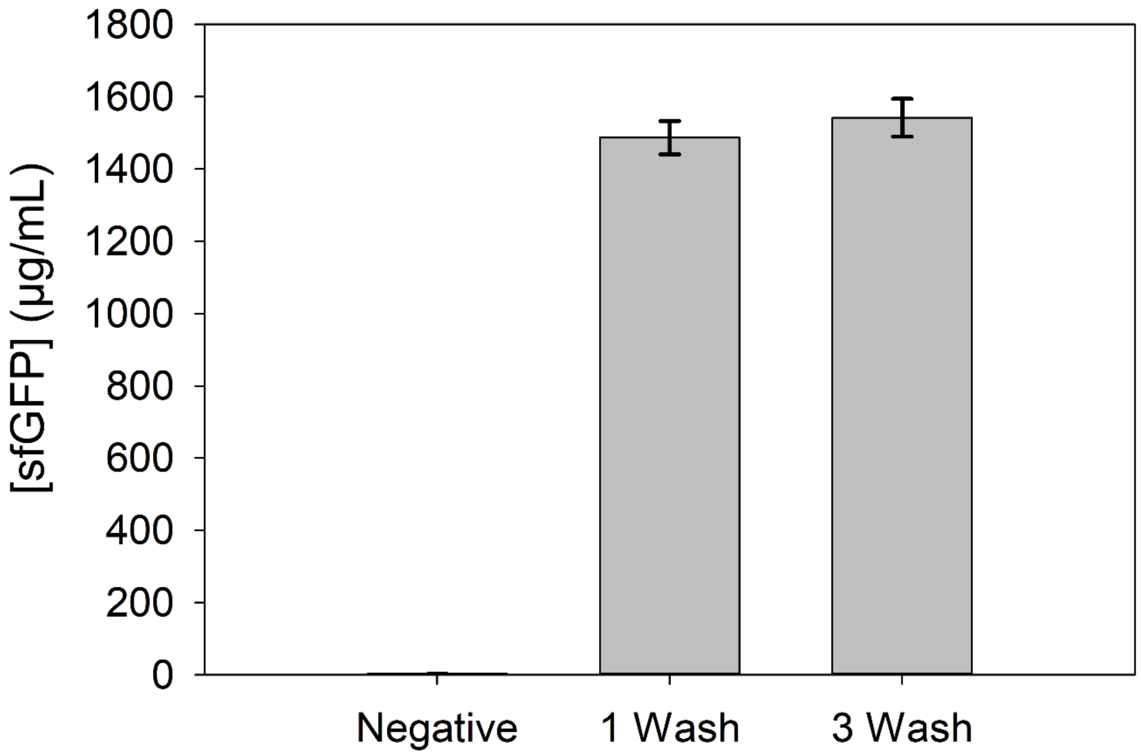


**Supplemental Figure 2**. Comparison of one wash versus three washes of cell pellets during harvesting of growths in 2x YTPG media for preparation of high yielding extracts. A single pellet was split in half and underwent 3 washes versus 1 wash and then underwent extract preparation. The resulting extracts underwent CFPS reactions in triplicate for sfGFP, and the resulting fluorescence was measured in triplicate. All error bars represent one standard deviation of the average of three independent reactions for each condition.

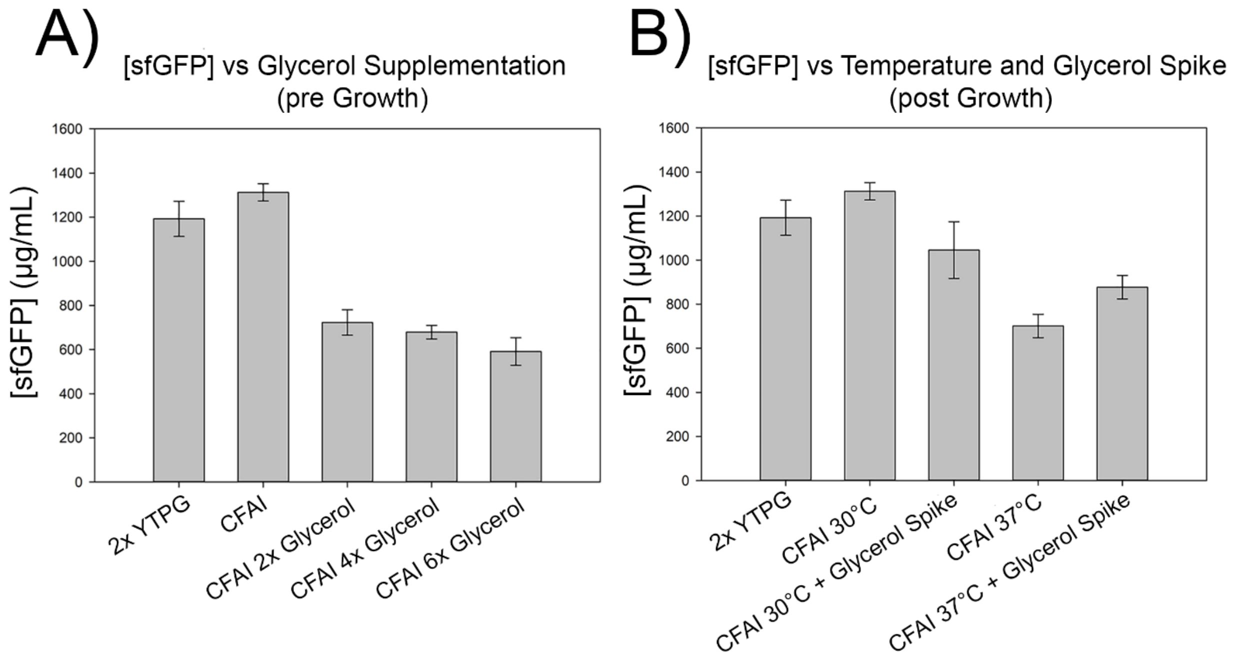


**Supplementary Figure 3**. Glycerol supplementations to CFAI media prior (A) and post overnight growth (B) resulted in decreased [sfGFP]. CFAI media formula underwent a 2x, 4x, and 6x titration of the 1x glycerol formula in panel A (6mL of 100% glycerol). CFAI overnight growths represented in panel B were grown at 30°C and at 37°C with one from each respective temperature undergoing supplementation with 6 mL of 100% glycerol one hour prior to harvest. Values represent averages across three independent reactions measured in triplicate. All error bars represent one standard deviation of the average of three independent reactions for each condition.

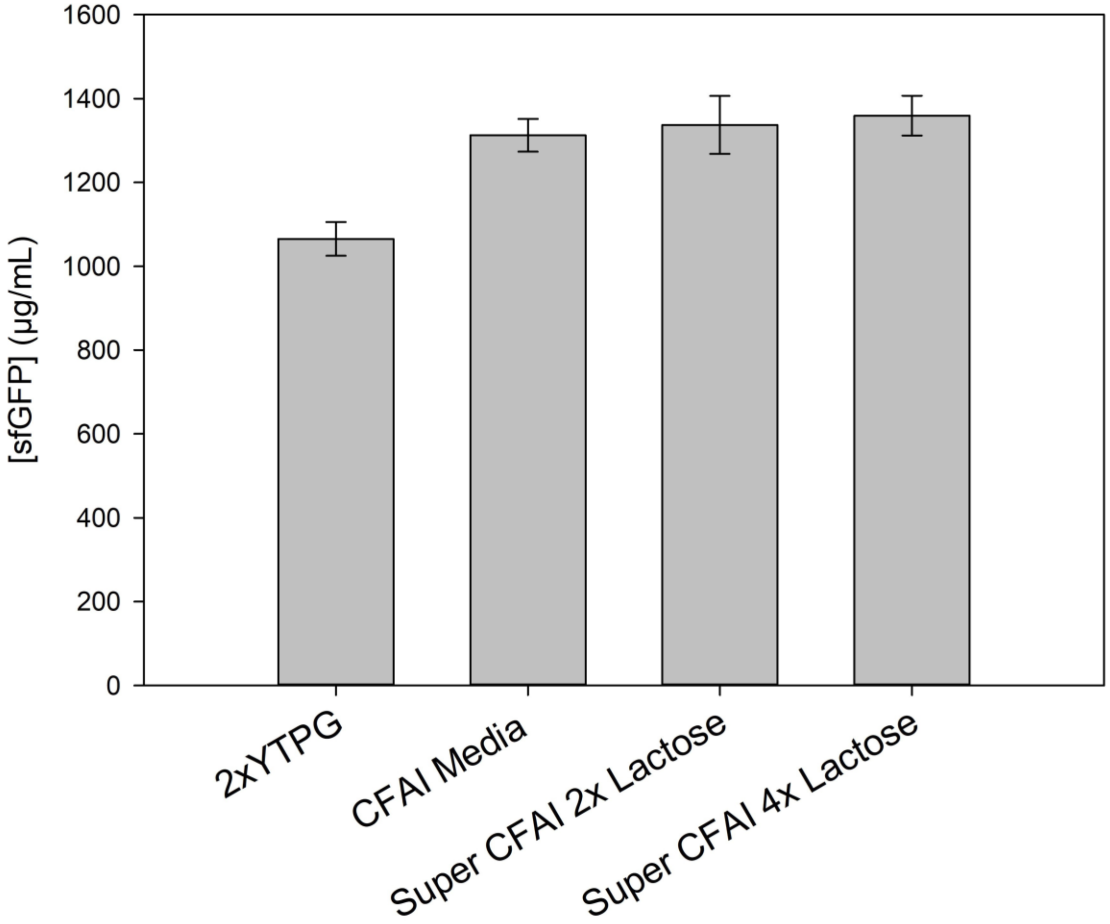


**Supplementary Figure 4**. Tryptone, yeast, and lactose supplementations to the CFAI media formula displayed no added boost to [sfGFP]. Recipes for various media types are located in Supplementary Table 1. Values represent averages across three independent reactions measured in triplicate. All error bars represent one standard deviation of the average of three independent reactions for each condition.

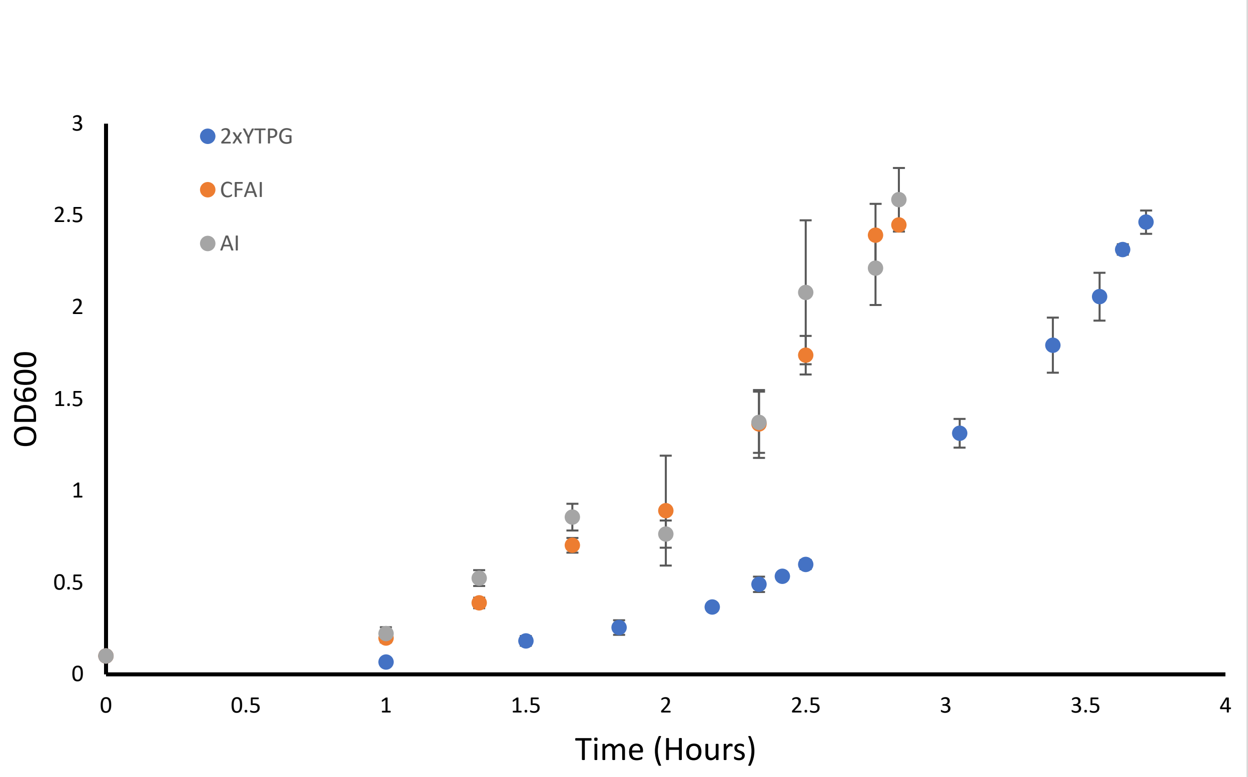


**Supplementary Figure 5**. Growth rates of various media conditions from $OD_{600}$ of 0.1 to 2.5 resulted in a significantly faster growth rate in autoinduction media (CFAI in orange; AI in grey) compared to 2x YTPG media (blue). Values are averages across growths performed in triplicate for each media type. All error bars represent one standard deviation of three independent growths for each condition.

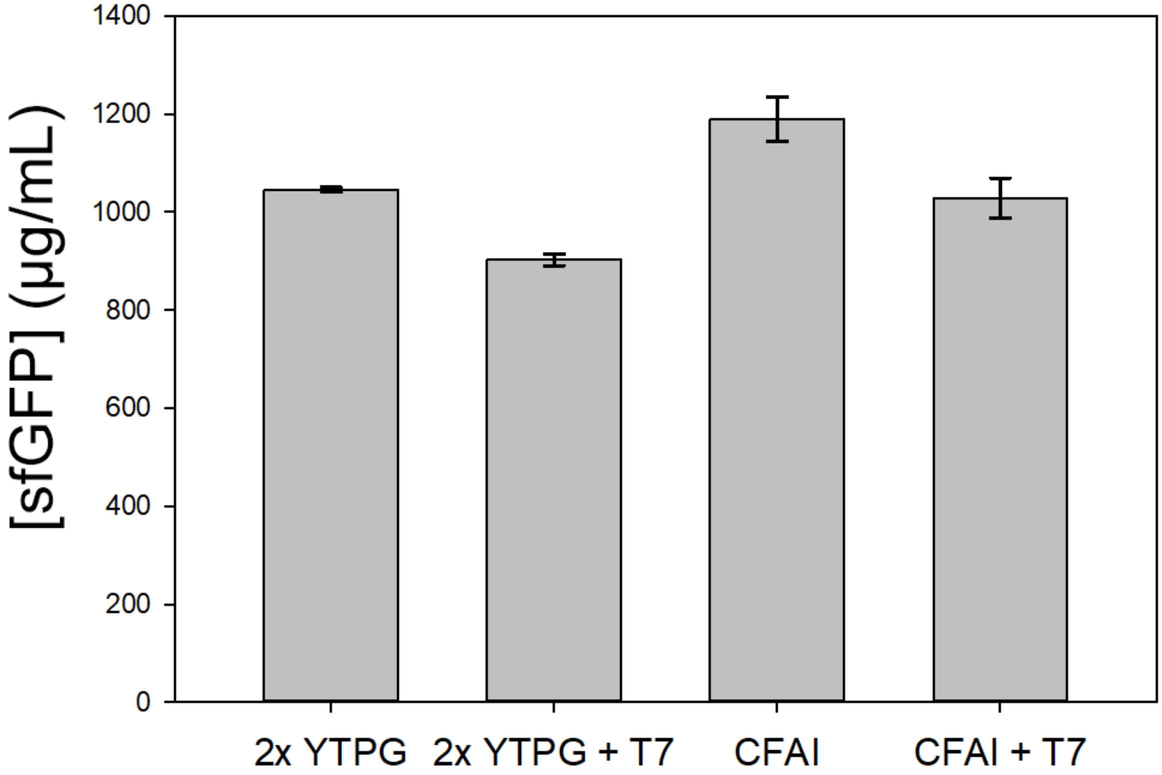


**Supplemental Figure 6**. Evaluation of T7 RNAP limitations in 2x YTPG and CFAI media. Extracts from 1 L growths of each media type were tested by cell-free protein synthesis reactions in triplicate and production of sfGFP was measured in triplicate for each reaction. [sfGFP] is not improved with exogenous addition of T7 RNAP across media types. All error bars represent one standard deviation of the average of three independent reactions for each condition.
